# RNA-seq variants reveal distinct patterns in the aging epitranscriptome: an in-depth analysis of age-matched Alzheimer’s Disease patients and a cognitively normal cohort

**DOI:** 10.64898/2026.02.14.705891

**Authors:** Katherine A. Jensen, Jennifer Weller, Kristen E. Funk

## Abstract

**Background:** Post-transcriptional modifications are those made to the RNA transcript, which can modulate RNA stability and function. Despite robust investigation of the genome, transcriptome, and proteome, little is known about post-transcriptional modifications during normal aging or Alzheimer’s disease (AD) pathogenesis. Several studies have shown epitranscriptomic changes in AD brains for certain modification types, establishing epitranscriptomic links to the disease; however, the complete set of post-transcriptional modifications have not been assessed in the context of AD. Furthermore, it is not understood which genes or pathways are under epitranscriptomic regulation, how conserved and sporadic modifications are distributed, or which conserved sites are differentially modified in diseased brains. Therefore, there is a need for a more complete analysis to describe the full landscape of the epitranscriptome in AD, helping to bridge the knowledge gap between post-transcriptional modifications and the molecular etiology of AD.

**Methods:** We designed and implemented a novel bioinformatics pipeline for complex epitranscriptome-wide analysis of potential RNA modification sites in sample-matched, whole-genome sequencing-filtered variant calls from RNA sequencing data. Using parametric and non-parametric tests, we tested differences in patterns for all detectable variant calls between postmortem brains of AD and cognitively normal, aged individuals.

**Results:** We identified 544 genes with hyper-modified transcripts in AD samples compared with cognitively normal controls, a notable observation being high enrichment of genes in the “Kaposi’s sarcoma-associated herpesvirus” pathway. We also identified patterns of recurring and sporadic modification sites that differed complementarily between disease and non-disease conditions. We found 17 genes (33 total sites) that were differentially modified between conditions including several sites found exclusively in the AD epitranscriptome.

**Conclusions:** These findings provide a more complete profile of the potential molecular underpinnings which differentiate AD brains from their non-diseased, aged counterparts and reveal patterns and modification sites which can be further investigated for how they contribute to the network of molecular interactions underlying AD. These elements are likely to be valuable candidates for investigations that aim to further the search for biomarkers and therapeutic targets.

## Background

Despite decades of research on neurodegeneration in aging populations, the molecular changes that underlie Alzheimer’s disease (AD) are still not fully understood. While a set of genetic markers has been defined for the low-occurrence Early Onset Familial Alzheimer’s Disease (EOFAD), no universal molecular underpinnings for the most commonly occurring Late Onset Alzheimer’s Disease (LOAD) have been identified (occurrence rate ∼ 90 – 95% of all AD cases) (1). Often, a diagnosis is not made until the cognitive capacity of a patient has drastically deteriorated; at which point, little can be done to mitigate progression of the disease. As average human lifespans increase, the prevalence of LOAD has risen exponentially, yet non-invasive early diagnostics and cost-effective interventions remain elusive (2,3). While significant disease-contributing factors have been identified from the genome (4–7), proteome (8), and transcriptome (9–13), a comprehensive ‘omics-level analysis of the epitranscriptome in aging and AD has not yet been reported. Given that neurons are post-mitotic cells, and thus, non-dividing, their epitranscriptomic regulation is likely to be one of the primary drivers of altered neuronal activity in aging and diseased brains.

Epitranscriptomic modifications are chemical changes made post-transcriptionally to the nucleotides of RNA transcripts. These changes include A-I and C-U editing, methylation, and pseudouridylation. Such alterations are dynamic, and can initiate adaptive responses to both internal (14,15) and external stimuli (16–18). By altering regulatory signals, the epitranscriptome modulates gene expression levels and alternatively spliced isoforms without changing the genomic sequence (12,14,19).

Some RNA modification types and positions are conserved, found consistently across groups, especially in highly conserved RNA structures such as transfer and ribosomal RNAs (20). Other modification types and sites arise more sporadically, occurring only in certain contexts and not uniformly across or within groups (21,22). Furthermore, the proportion of modified transcripts out of all transcripts present for an expressed gene can vary greatly, altering the level of proteins and their isoforms that are present to perform a range of tasks in the cell for a particular environment at a given time (23–25). Therefore, despite the ubiquity of post-transcriptional modifications, two distinct, independent profiles can describe an individual’s epitranscriptome: one profile, derived from binary counts in epitranscriptomic data, represents the presence or absence of a modification and is useful for determining modification load between groups; the other, calculated from the proportion of transcripts modified, represents the stoichiometry of the modifications that are present, identifying the differential modification occupancy for a given transcript, sample, or group.

While many experimental methods exist to reliably detect the presence of epitranscriptomic modifications in research, they are less suitable for transcriptome-wide assessments as they often only interrogate for a single modification-type at a time and can produce low-resolution results if multiple modifications are within close proximity to one another (26,27). While it is not yet possible to identify all modification types in a high-throughput manner, it is possible to find sites that have reference-mismatched nucleotide calls from RNA sequencing (RNA-seq): these mismatches are a result of modification-induced errors that are introduced during the cDNA library preparation (28–31). Therefore, several computational tools and algorithms for epitranscriptomic analyses of high-throughput RNA-sequencing datasets have been developed (32). Such tools demonstrate that it is possible to identify post-transcriptional patterns from RNA-seq (33); however, most of these tools focus primarily on A-I editing and rely on A-I edit databases. None of the established tools inclusively assess all filtered variant call-types nor provide quantitative analysis of recurring versus sporadic modifications per site, modification stoichiometry, and differential modification usage by gene, sample, and condition.

The purpose of this study was to establish an in-depth analysis of variations from RNA-seq data to better define the molecular drivers and functional outcomes of such epitranscriptomic changes. To address the current gaps in knowledge, we created a custom epitranscriptome analysis pipeline which builds on previously established approaches and adds novel algorithms created for the purpose of this study. We aimed to a.) establish global modification differences between AD and age-matched cognitively normal (CN) brains, b.) assess the distribution of recurring and sporadic modifications within and between conditions, c.) interrogate differential modification load and stoichiometry, and d.) determine pathways under epitranscriptomic regulation in AD and CN brains. Our results show that epitranscriptomic landscapes differ between AD and CN brain samples in global modification site numbers and/or stoichiometry, specific transcripts subject to such modifications, and the nucleotide positions modified. From our most significant results, we were able to identify connections between the epitranscriptome, viral infection, neuroinflammation, and AD. These results provide information regarding molecular changes that may drive AD related neuropathology as a consequence of modified RNA transcripts or through shared mechanistic pathways. We anticipate that these results will provide insight into the molecular etiology underlying the neuropathology and pathophysiology of AD.

## Methods

### Data retrieval, quality control, and filtering

Paired-end whole-genome sequencing (WGS) and RNA-seq data produced from the Illumina HiSeq and NovaSeq platforms (34), respectively, were derived from ‘bulk brain tissue’ samples of postmortem brains from the ROSMAP datasets. Data was securely uploaded to the UNC-Charlotte HPC cluster via the Synapse python client (35). Datasets analyzed were WGS: syn20190457, syn20068561 and, RNA-seq: syn3388564. Specifically, patient data that contained matched RNA-seq and WGS data were retained for analyses. De Jager et al provide an in-depth description of the sample processing and data generation protocol (36). According to the synapse webpage: “for RNA-seq data, samples were extracted using Qiagen’s miRNeasy mini kit (cat. no. 217004) and the RNase free DNase Set (cat. no. 79254) and quantified by Nanodrop UV and RNA quality was evaluated by Agilent Bioanalyzer. WGS sequencing libraries were prepared using the KAPA Hyper Library Preparation Kit in accordance with the manufacturer’s instructions.” Patient metadata is described in **Table 1**. Samples were divided into two conditions, based on clinical diagnosis of cognitive status (*dcfdx*) where 1 == No Cognitive Impairment (Cognitively Normal; CN) and 4 == Alzheimer’s (disease) dementia with no other contributing factors to cognitive impairment (AD). Some individuals assigned to AD-level cognitive status were not recorded as having received a clinical diagnosis of AD dementia (*age_first_ad_dx*) and these samples were removed from our analysis. Since the metadata identifies 90+ ages into a single age category, age counts were grouped according to < 90 or 90+ and a chi-square test of independence (Python (37); *chi2_contingency()* function from the *scipy.stats* module) was used to check for age-related biases between conditions. Data were filtered and processed as outlined in **Figure 1**.

**Figure 1.**
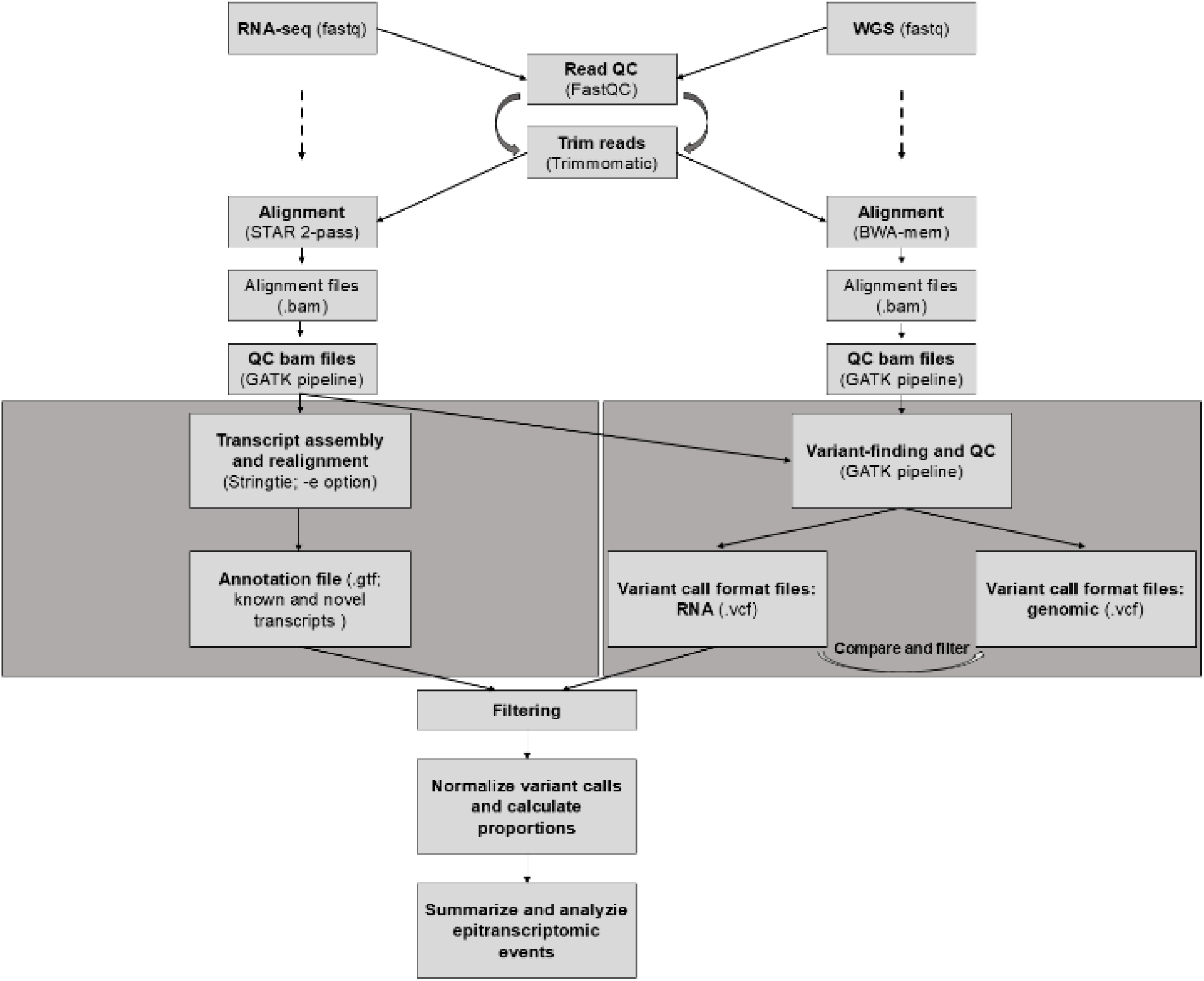
Data processing workflow. RNA-seq and WGS raw FASTQ files were trimmed and filtered for quality. Both datatypes were aligned to the reference human transcriptome and genome, respectively, and the output files were filtered again for quality. Stringtie was used to identify transcripts most likely present in the dataset, along with calculated FPKM and other values for each transcript per sample. Frequently cited GATK variant-calling pipelines were applied to extrapolate the presence and position of RNA-specific variants. Variant-called RNA data was further processed for downstream analyses.

**Table 1.** Demographics of Participant Samples.

|  |  | CN |  | AD |  |
| --- | --- | --- | --- | --- | --- |
| Age |  | 73.8 – 89.9 | 90+ | 80.3 - 89.9 | 90+ |
| Sex |  | 13 (43%) / 5 (17%) | 9 (30%) / 3 (10%) | 10 (34%) / 2 (7%) | 14 (48%) / 3 (10%) |
| Female/Male |  |  |  |  |  |
| **APOE** | 2/2 | 0 | 0 | 0 | 0 |
|  | 2/3 | 4 | 2 | 1 | 1 |
|  | 2/4 | 0 | 0 | 0 | 0 |
|  | 3/3 | 12 | 8 | 4 | 11 |
|  | 3/4 | 1 | 2 | 6 | 4 |
|  | 4/4 | 1 | 0 | 1 | 0 |
| **Braak** | I - II | 3 | 4 | 1 | 1 |
|  | III - IV | 12 | 6 | 6 | 4 |
|  | V - VI | 3 | 2 | 5 | 12 |
| **Cerad** | 1 - 2 | 18 | 12 | 0 | 0 |
|  | 3 - 4 | 0 | 0 | 12 | 17 |

*Note.* APOE: Status of Apolipoprotein E genotype. Braak: neurofibrillary tangle stages, higher stages indicate greater tangle burden. Cerad: Amyloid- $\beta$ plaque score, higher scores indicate greater plaque density. APOE assignment is missing for (n=1) AD brain sample.

An anaconda v3 environment was created to manage tools and modules needed for the data processing, filtering, and analysis pipelines. Raw sequencing reads (FASTQ) for RNA-seq and WGS dataset-types were first run through a read quality control step. FastQC (38) was used for validation of read quality and Trimmomatic v0.39 was utilized for removal of adapter/index sequences and/or low-quality reads, with use of various flags and parameters depending on the quality score patterns provided by FastQC and the length of the reads in the data subset. BBmap’s v38 (39) *filter-by-tile* script (with ‘lowqualityonly=t’ and ‘overwrite=t’) was run to remove low-quality tile data.

### Quantification and qualification of transcripts

In order to retain potentially novel transcripts and to identify baseline expression of all transcripts, we captured all but the shortest possible transcripts (minimum length 50 bases), RNA-seq FASTQ files filtered by quality and length (50nt minimum) were analyzed using Stringtie (40) v2.2.0+ in a 3-step process: 1.) Stringtie was applied to each sample (’--rf -t -p - B’), 2.) assembly information from the first step was merged (’--mergè) into an updated transcriptome, and 3.) Stringtie was applied again, this time using the updated transcriptome information from the second step (’--rf -m 50 -t -p -B -è). Along with the Stringtie output per individual, a General Transfer Format (GTF) file containing annotations for all transcripts identified by the program was created and saved for subsequent analyses. Using python’s v3.0+ *gfftutils* library, a FASTA file of the novel transcripts was created from the Stringtie-generated GTF file and merged with GENCODE’s GRCh38 reference FASTA for use in next steps.

### Alignment, filtering, and variant-calling

To define a set of post-transcriptional modification sites that could be compared between disease and non-disease states, sequencing data was aligned to a reference and filtered on alignment quality and read depth. The data were not haplophased: therefore, “allele” is used here to define the genomic site to which a nucleotide in the RNA-seq reads mapped, and “alternate allele” denotes the presence of a nucleotide different from that in the reference transcript.

Alignment to *H. sapiens* reference genome GRCh38 (41) was the first step for variant finding. RNA sequencing reads were aligned using STAR (42) v2.7 two-pass mode, which incorporates novel junctions found in the data into the reference for a sample-customized alignment (with ‘--twopassMode Basic’), and WGS data was aligned with BWA-mem (43) v0.7+ (default parameters). Using the GATK v4.2.0 best practice pipelines for small variant discovery (44,45), we filtered and processed the RNA-seq and WGS alignment files. First, FASTQ files were converted to unmapped BAM files with *FastqToSam*/*Fastq2Sam* (’-DS’ for RNA-seq “Description” field in addition to required flags and default parameters) and the unmapped BAMs were merged with aligned BAM files with *MergeBamAlignment* (default parameters).

Merged BAM files were then run through a series of quality-control steps, sorting and marking duplicates with a combination of samtools v1.11 (46) and tools from the GATK Picard suite: *SortSam* (’-SO queryname coordinatè), *MarkDuplicates* (default parameters), and *BuildBamIndex* (default parameters). R v4.0.3 (47) was loaded for GATK tools that require R for proper functionality. Additional tools run on the RNA-seq data at this step also included *SplitNCigarReads* (default parameters) to reformat spliced reads for compatibility with downstream analysis in the pipeline and *AddOrReplaceReadGroups* (’--RGSM –RGID –RGDS --RGPL --RGLB’) to store metadata not included by default in the BAM files produced by STAR. To calculate differences between observed error rates and the sequencer’s reported Q-scores, *BaseRecalibrator* (default parameters) was applied, using known sites from hg38 dbsnp138 (48). *ApplyBQSR* (default parameters) was then used to adjust read base quality estimates from the *BaseRecalibrator* output. After this point, the RNA-seq and WGS datasets were analyzed independently.

In the final variant finding steps for the RNA-seq dataset, *AnalyzeCovariates* (default parameters) was used to generate plots allowing assessment of the effect of base-recalibration processes on the quality scores, variants were called using *HaplotypeCaller* (’--output-mode EMIT_ALL_CONFIDENT_SITES’) and then filtered with the *VariantFiltration* tool (’--filter-name ‘qual_filt’ --filter-expression ‘QUAL < 25.0’ --filter-name ‘qual_depth_filt’ --filter-expression ‘QD < 10.0’ --filter-name ‘depth_filt’ --filter-expression ‘DP < 20’ ‘). Filtered variant call format (VCF) files were used in downstream analyses.

The final steps of variant-finding for the WGS dataset applied the *Mutect2* tool in “tumor-only” mode (’-tumor --max-mnp-distance 0 ‘) on the cognitively normal dataset, since a panel of normals (PONs) is required in order to call SNVs and indels. There is a publicly accessible PONs available through GATK, but the panel was created from a cohort of relatively young, presumably healthy individuals. To reduce age-related biases, a PONs was created from the aged study cohort’s derived variant call information from Mutect2’s “tumor-only” mode as follows: the *GenomicsDBImport* tool (default parameters) was utilized to create a database from the CN variant calls; the *SelectVariants* tool (default parameters) was used to create an exported variant-call format file from the database; the database-derived VCF file was input to *CreatePanelOfNormals* tool (default parameters) to create the reference PONs. Variant-finding was then conducted on the samples with *Mutect2* (’ --max-mnp-distance 0 --panel-of-normals PONs ‘). Finally, VCF files created by *Mutect2* for each sample were run through *GetPileupSummaries* (default parameters) to collect pileup summaries at common variant sites and estimate allele fractions in the sample, then *CalculateContamination* (default parameters) was run to create a contamination table for *FilterMutectCalls* (’ --contamination-table ‘) for a final filtration of cross-sample contamination and possible sequencing artifacts. The filtered VCF files were then used in downstream analyses.

### Elucidating RNA-specific modifications

Genomic variants in transcribed regions are likewise observed in transcript sequences, so a custom python script was used to compare RNA variants against WGS variants and filter out matched sites. Only sites with a variant call in the RNA but not in the genomic DNA for each individual were retained for downstream analysis.

### Comparison of modifications between AD and CN groups

With python’s *sqlite3,* a working database was created and tables were populated with information from 1.) cDNA and lncRNA sequences (GENCODE v42; GRCh38), 2.) RNA VCF files created from STAR alignment, and 3.) a GTF file (GRCh38 annotations merged with Stringtie annotation output). Queries across the database enabled output suitable for both global-and local-level comparative analysis pipelines, including statistical tests and p-value filtering of modified genes and variant call-types as illustrated in **Figure 2**.

**Figure 2.**
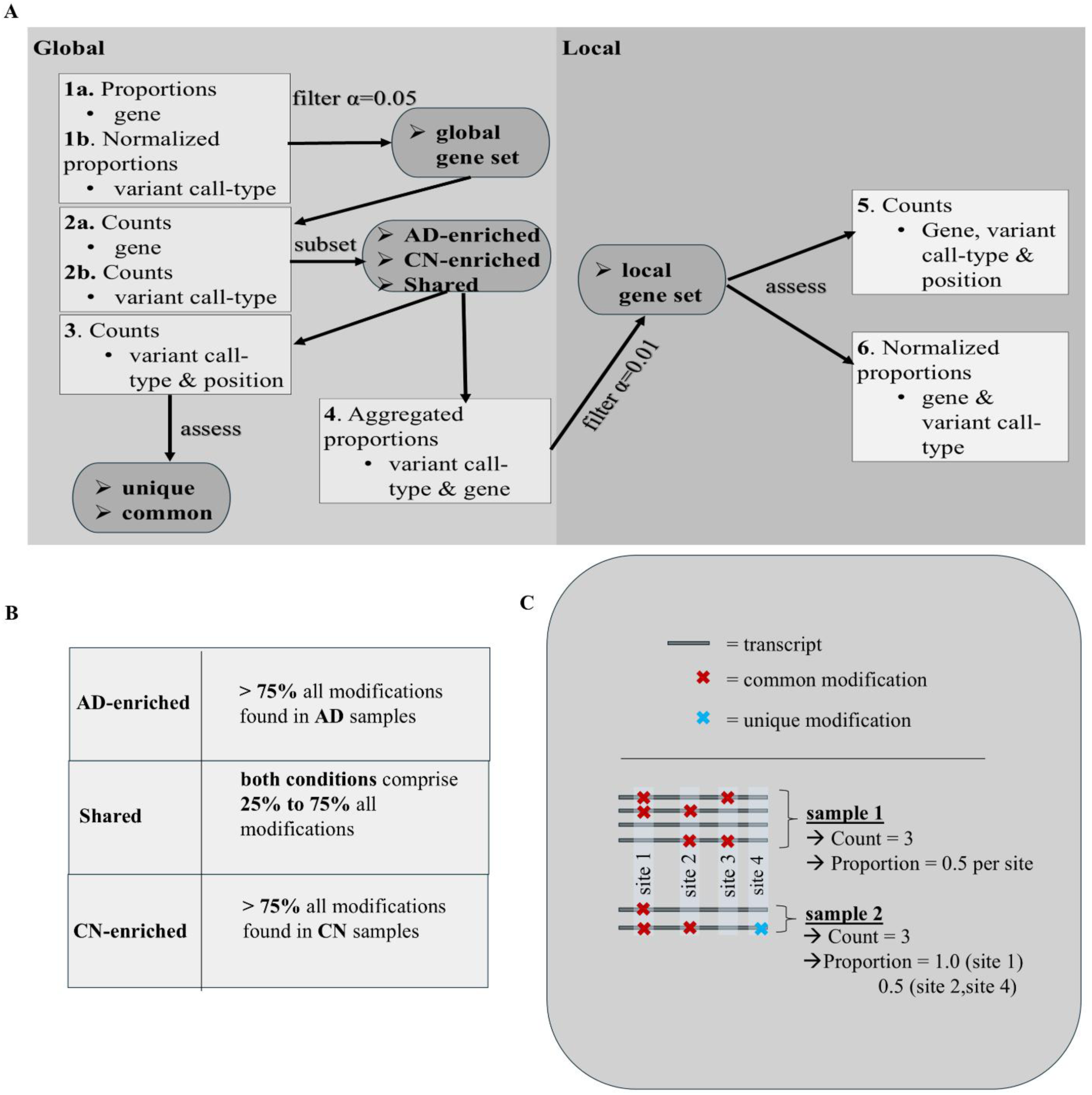
Data analysis pipeline. **A)** Diagram of the global to local analysis pipeline for identification of epitranscriptomic modification patterns in aging and AD. Steps 1 and 6 p-values were derived from a Mann-Whitney-U test. A two-sample independent t-test produced the p-values for Step 4 filtering. The AD-enriched gene subset from Step 2 was further assessed for pathway enrichment. **B)** Criteria for sub setting genes into AD-enriched, CN-enriched, or Shared categories to determine distribution of modification burden at the global level. **C)** Schematic of the differences in counts vs proportions for assessment and filtering of modifications. Proportions were calculated per site, where the number of modifications (allele depth; ad) was divided by the number of expressed transcripts (supporting read depth; dp). Counts indicate the total number of modified sites identified for a sample. Unique modifications are those found only once in the entire sample space. *Note*: Per-position proportions were normalized by length of gene for gene-to-gene comparisons: (ad/dp) * (1/length). Aggregated proportions were calculated as the sum of allele depths divided by the sum of supporting read depths for a sample, without normalization, for within gene comparisons.

### Assessing global variant call-type usage between conditions

Starting with genes for which a minimum of one RNA variant call count was identified in all samples of at least one condition, we analyzed epitranscriptomic modification profiles at the global level. Proportions were calculated per site, for within-gene comparisons, by dividing allele depth (*ad*) by the number of supporting reads (*dp*):

(*ad/dp*)

To reduce gene length biases, when comparing features across genes, proportions of modification level at each position were normalized by length, where per-site proportions were multiplied by a position weight (*pw*), calculated as 1/length_transcript_:

(*ad/dp*) * (*pw*)

We used a Mann-Whitney-U test with Python’s *scipy.stats.mannwhitneyu()* to evaluate the differences in modification stoichiometry between conditions for each gene (within genes) and for each variant call-type (across genes). From the within-gene tests, only genes that had a significant difference (α=0.05) in total modification proportions between conditions were retained in a global gene set for the next steps (**Step 1a; Figure 2A**). P-values, derived from comparisons between conditions of normalized per-site proportions (**Step 1b; Figure 2A**) across all genes, were used to determine global differences in modification stoichiometry by variant call-type.

Using the collection of modified sites from genes in the global set, overall modification burden per variant call-type was assessed using sums of modified sites for each condition as shown in **Step 2b** of **Figure 2A**. Python’s *sqlite3()* was used to query the database with SELECT COUNT(*) summing each variant call-type, per sample per condition, across all genes. A CSV file was created from these counts and R *ggplot2()* was used to create a boxplot. The boxplot was further annotated with *p*-value significance levels at 0.001, 0.01, and 0.05, as were reported from the stoichiometric comparisons for each variant call-type (**Step 1b**; **Figure 2A**).

### Assigning filtered genes to condition-enriched subsets

As shown in **Step 2a** of **Figure 2A**, we determined which genes from the global set were hyper-modified in one condition over another and assigned each gene to one of three possible global condition-enriched subsets. Subset assignment was based on percent of sites modified in the gene for all samples in a condition out of all positions modified across both conditions.

To create these assignments, the number of modified sites per sample, were summed (counts) across all samples in the AD and CN datasets. Then, the sum for each condition was divided by the total modified sites found in the gene and the resulting quotient multiplied by 100. As illustrated in **Figure 2B**, the resulting percentages were used to assign each gene to one of 3 global-distribution subsets: AD-enriched, CN-enriched, or Shared. “AD-enriched” and “CN-enriched” distributions were assigned when 75% - 100% of all modifications for a gene were identified in the same condition, and the “Shared” distribution was assigned when variant counts in both conditions constituted 26 – 74% of all variants for that gene.

### Testing differences in the distributions of sporadic and recurring modifications between conditions

Binary counts of uniquely and commonly modified positions were analyzed to determine the distribution of sporadic and recurring modifications between conditions as shown in **Step3 Figure 2A** and illustrated in **Figure 2C**. A ‘SELECT DISTINCT COUNT’ query of the database table containing sample information was used to sum modification sites for each variant call-type.

Counts were summed for each condition, variant call-type, and condition-enriched gene subset. From the summed counts, distributions of sporadic modifications (unique; identified in only one sample across both conditions) and recurring modifications (common; identified in more than one sample of either or both conditions) were compared using a two-sided Fisher’s Exact Test with the *fisher_exact()* function from Python’s *scipy.stats* module. Log odds ratio (OR) outcomes from the test were subject to Haldane-Anscombe correction for zeros, then mapped as a forest plot with the *plt()* function from Python’s *matplotlib.pyplot* module.

**Additional File 1** contains the count sums as input for the Fisher Exact Test and the test odds ratio and p-value output which were used as input for the plotting function.

### Pathway analysis of AD-enriched genes

To better understand which pathways are most likely to be affected by post-transcriptional modifications in diseased brains, the AD-enriched subset of genes was submitted to Enrichr-KG (49) and analyzed by the KEGG 2021 Human library for pathway information using the KEGG and GO database options.

### Filtering and assessing modifications at the local level

To determine the overall modification stoichiometry by variant call-type per gene, as shown in **Step 4** of **Figure 2A**, proportions within each gene were aggregated per variant call-type by dividing the sum of the total allele depth counts with the sum of the total supporting read depth counts for each modified site from all samples. The difference in overall modification stoichiometry by variant call-type between the conditions was tested with a two-sample independent t-test on the aggregated modification proportions for each gene. Genes with a significant difference (α=0.01) in at least one variant call-type between the conditions were assessed at the local level. As shown in **Step 5** of **Figure 2A**, we determined how many samples had each modified site of the variant call-type(s) determined to be stoichiometrically different in genes from the local gene set by summing and comparing the modified site counts. A ‘SELECT DISTINCT COUNT(DISTINCT)’ query grouped by gene, variant call-type and condition was used. Finally, a more sensitive method was used to determine the local modification stoichiometry for each variant call-type using a Mann-Whitney U test on normalized proportions for each sample (**Step 6; Figure 2A**), and the average proportions were compared with a boolean operation to determine which condition had the highest average proportions. This set of genes and positions were used to create local level profiles with only those that showed either a) 50% or more of the samples in a condition were modified at a site while no samples in the other condition contained the modification, indicating per-site modification load (see **Step 5 Figure 2A**), or b) a significant difference (α = 0.05) in stoichiometry between conditions for all sites in a gene with the variant call-type as determined by Mann-Whitney U test (see **Step 6 Figure 2A**), extrapolating the differential modification occupancy by variant call-type. The Pathway Commons Protein-Protein Interactions database (50) and the literature were searched for additional pathway information involving these genes.

## Results

### Samples are age-matched

AD and CN samples from the ROSMAP bulk-brain dataset were filtered to include only sample sets with both RNA-seq and WGS data, as described in the methods. After this filtering step, 59 samples remained: CN (n=30) and AD (n=29). To determine if an age bias was present in the retained samples, we performed a Chi-square test for independence, which determined no statistically significant difference in age distributions between conditions (*Χ^2^* = 1.37; *P*-value = 0.24; *df* = 1) and therefore it is not expected that age differences confound interpretation of results. Sample information included in the metadata is further described in **Table 1**.

### Variant calls indicative of epitranscriptomic modifications are increased in AD

To first establish the background rate of post-transcriptional modifications in the dataset, all variant call-types were assessed equally and counts of modification sites by variant call-type were summarized and filtered as shown in **Figure 1**, resulting in 22,574 distinct genes with modification rates at or above a minimum threshold of one site each for all samples in at least one condition. When global differences in modification stoichiometry were tested between AD and CN conditions for modification sites in each gene as shown in **Step 1a Figure 2A**, 679 genes were found to be significantly different between the conditions and comprised the global gene set. Sums of variant call-types for each sample from genes in the global set, as shown in **Step 2b Figure 2A** revealed 62,683 total variant calls.

When modification counts for all samples were summed by each variant call-type across all genes in the global set of 679 genes, we found that all variant call-types appeared in the AD group at a higher frequency than in the CN group, as indicated in **Figure 3**.

**Figure 3.**
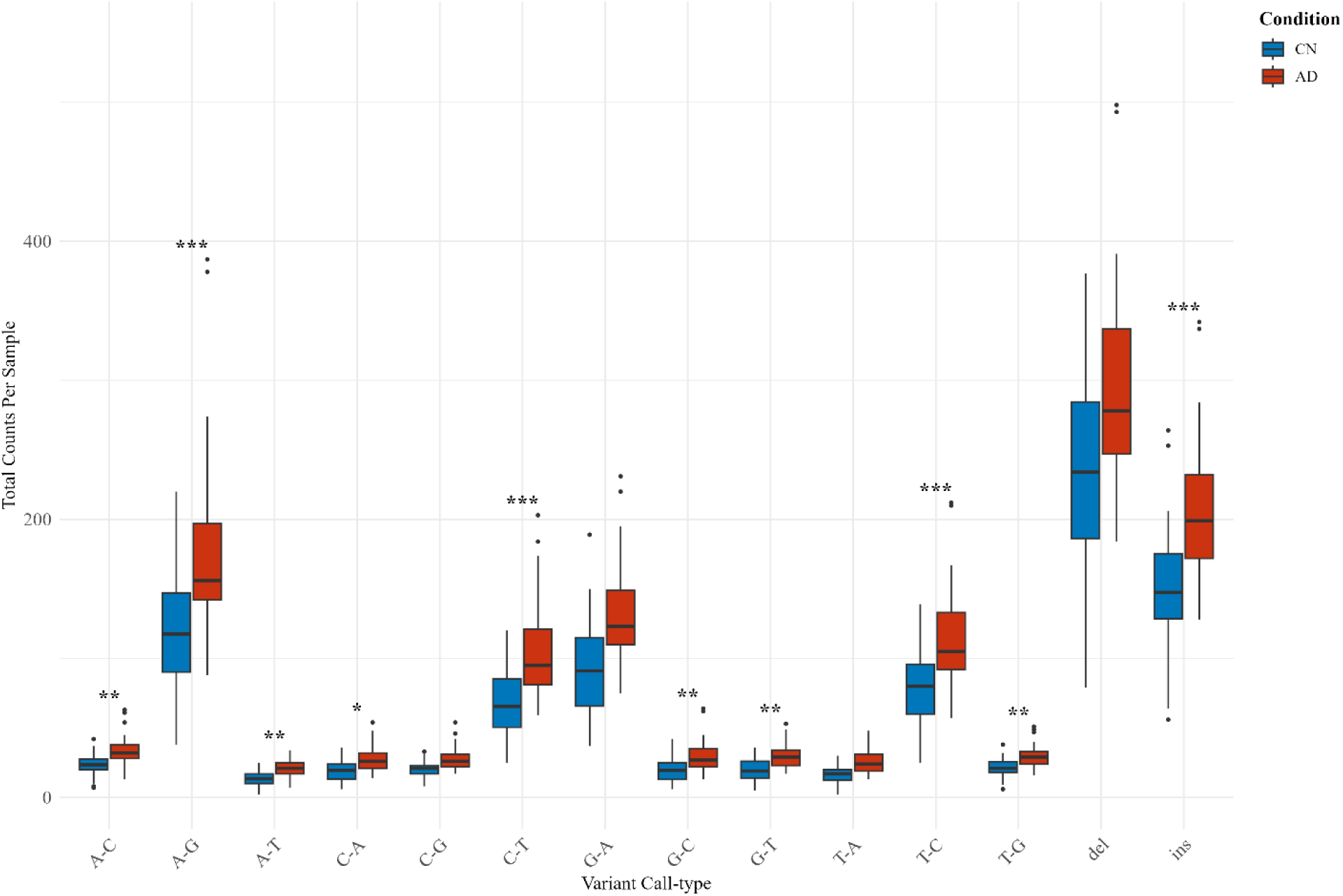
Frequency of RNA variants in bulk brain CN versus AD samples. Raw counts of modified positions across all samples and 679 genes by variant call-type. Data are shown as mean +/- standard deviation with outliers indicated by individual points. Statistical significance was calculated from Mann-Whitney-U test between normalized proportions of each sample in both conditions. *, *P* value < 0.05; **, *P* value < 0.01; ***, *P* value <0.001.

For both conditions, indels constituted the largest variant call-type group, with deletions occurring more frequently than insertions. The variant call-types that followed indels in order of frequency level, having at least 2,000 counts across all samples in both conditions, were as follows: A-G, G-A, C-T, and T-C. Across all genes in the global set, A-G, C-T, T-C, and insertions (ins), were significantly different between the two conditions at α=0.001; A-C, T-G, G-T, A-T, and G-C were statistically different at α=0.01; C-A variant calls were significantly different at α=0.05; C-G, deletions (del), G-A, and T-A were not significantly different between the two conditions.

These results show significant differences in modification stoichiometry of most variant call-types between the two conditions and global hyper-editing in the brains of AD patients, establishing the background of variant call-type distributions in the data.

### Transcripts for a subset of 631 genes were differentially modified between AD and CN

To specify the distribution of overall modification burden in AD and CN samples for each gene in the global gene set (**Step 1a; Figure 2A**), three condition-enriched subsets were created as shown in **Step 2a; Figure 2A** and detailed in **Figure 2B**. Assignment of a gene to one of three categories was based on comparisons of percent of total modification sites per condition, out of total counts recorded for the gene, resulting in 544 AD-enriched genes, 87 CN-enriched genes, and 48 Shared genes. These observations indicate that not all genes were modified equally, and distinguish those genes modified more often in either AD or CN brains.

### AD brains carry the majority of sporadic modifications in AD-enriched genes

To determine whether the distribution of sporadic and recurring modification sites differed between conditions, we analyzed each position by variant call-type across the condition-enriched gene subsets (**Step 3; Figure 2A**). Modified positions appearing only in a single sample across both conditions were classified as ‘unique,’ representing sporadic modifications. Modified positions appearing in more than one sample, whether in AD samples, CN samples, or both, were classified as ‘common,’ representing recurring modifications.

We compared the distribution of these counts in each condition using a two-sided Fisher’s exact test, followed by Haldane-Anscombe correction. The results demonstrated that unique modifications for nearly all variant call-types were statistically more likely to occur in the AD-enriched genes within AD samples than in CN samples. This trend was statistically significant except for C-A, T-G, A-C, and T-A, which did not reach the *p* < 0.05 threshold. Conversely, only insertions (INS), deletions (DEL), A-G and C-T were significantly more likely to occur in CN-enriched genes of CN samples than AD samples. **Figure 4** shows the distribution of log-odds ratio (LOR) confidence intervals and p-values for each variant call-type in both condition-enriched gene subsets. Together, these results show clear differences in recurring and sporadic modification distributions by variant call-type, in relation to sample condition and condition-enrichment of genes: AD-enriched genes contained the most sporadic modifications in AD samples while CN-enriched genes contained the majority of recurring modifications in CN samples.

**Figure 4.**
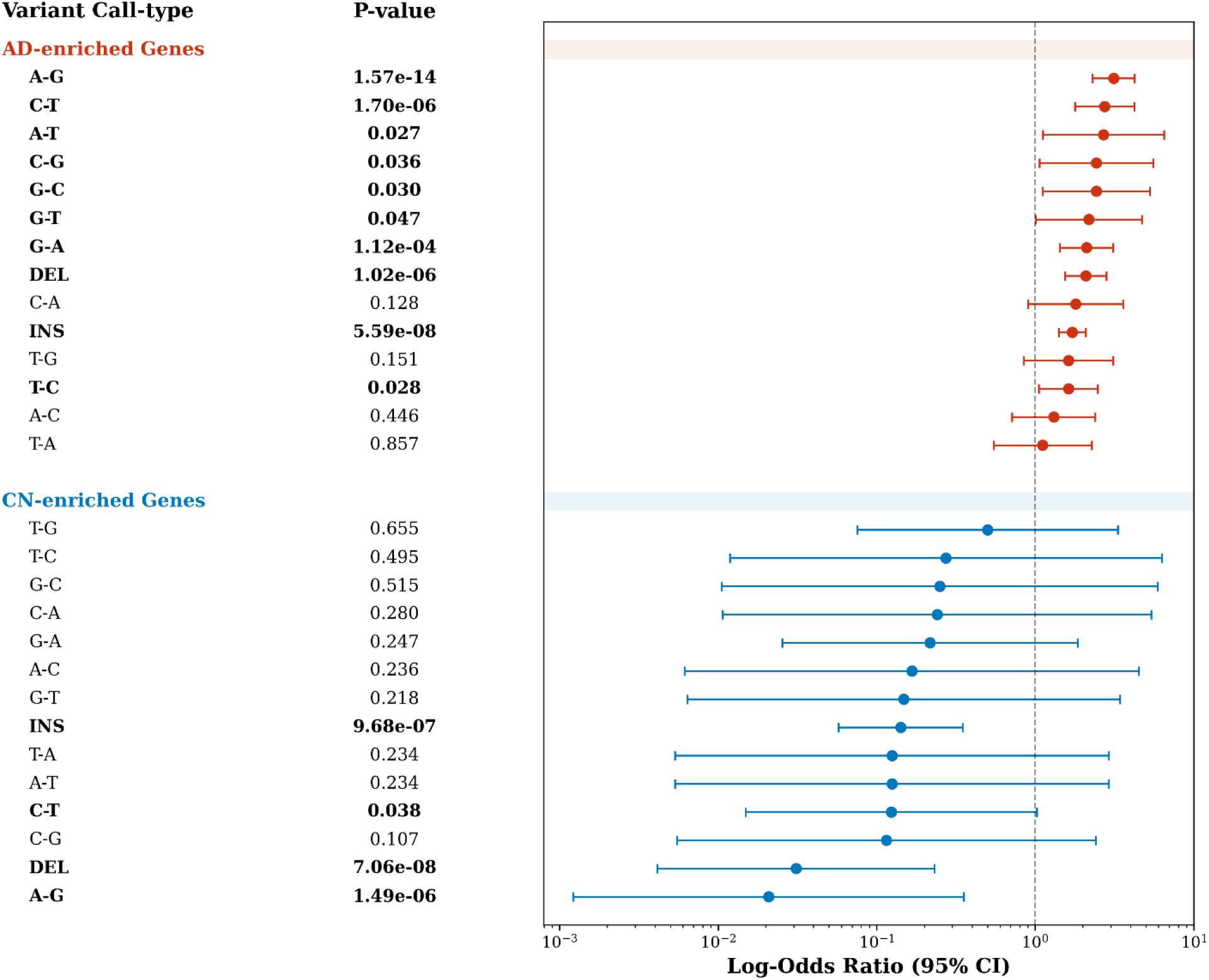
Forest plot of Log-Odds Ratio distributions for unique modifications in AD-enriched and CN-enriched genes. The central point of each line indicates the log-odds ratio (LOR) of unique modification likelihood, and the whiskers indicate the 95% confidence interval (CI). Red lines indicate AD-enriched genes and blue lines indicate CN-enriched genes. LOR < 0 indicates distribution in CN samples and LOR < 0 indicates distribution in AD samples. Significant association p-values are listed in bold type.

### AD-enriched genes are associated with processes governing viral infection response, cellular growth control, and protein processing

Before the final step of global-level analysis, AD-enriched genes were analyzed for gene pathway enrichment to determine which processes were associated with transcripts hyper-modified in AD brains. To investigate the pathway networks of genes influenced by higher levels of post-transcriptional modification in AD, all 544 of the AD-enriched genes were submitted to the Enrichr-KG interface. Results show the clustering networks and specific pathways that were over-represented in the “AD-enriched” gene set, which included “protein processing in the endoplasmic reticulum,” “renal cell carcinoma,” “chronic myeloid leukemia,” “Kaposi sarcoma-associated herpesvirus infection,” and “neurotrophin signaling” (**Figure 5**). While there is overlap among genes within the distinct the pathways, the “protein processing in endoplasmic reticulum” pathway had the most significant enrichment.

**Figure 5.**
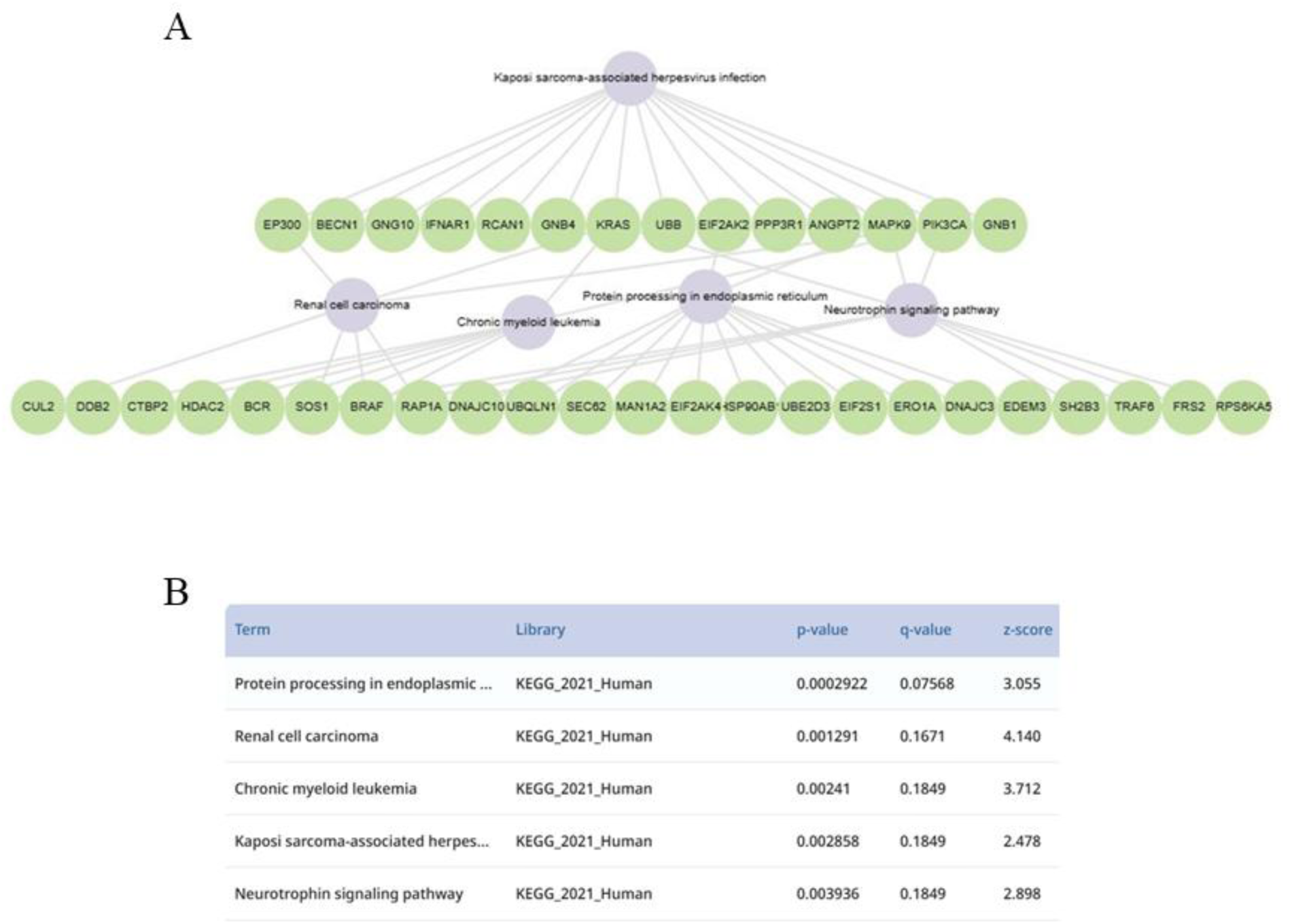
Enrichr-KG pathway analysis results for AD-enriched genes. **A)** Network visualization of the top enriched KEGG pathways and their associated genes from the AD-enriched subset. Purple nodes represent specific biological pathways, and green nodes represent the genes that contribute to those terms. Edges indicate the involvement of a gene within a particular pathway, highlighting significant gene-pathway overlaps. **B)** Summary table of the top five enriched pathways identified using the KEGG_2021_Human library. Pathways are ranked by their p-value, with additional metrics provided for q-value and z-score, indicating the statistical significance and degree of enrichment for each biological term.

### Distinct modification patterns differentiate AD brains from CN brains

Interrogating local patterns of each gene, position, variant call-type, and condition, we assessed epitranscriptome profiles at greater resolution than in the global analysis. From statistical testing of aggregated proportions, to determine the global modification stoichiometry by variant call-type for each of the 679 global subset of genes, 582 genes were identified as having a significant difference in at least one variant call-type (α = 0.05) between the conditions. From the set of 582 genes, only the (gene, variant call-type) pairs that were found to be significantly different between AD and CN at α= 0.01 were retained, resulting in 24 genes and 72 modified sites. When the retained set was filtered on counts and differential proportion p-values, a set of 17 genes and 33 modification sites that showed the greatest local differences in either modification load (**Table 2**) or stoichiometry (**Table 3**) were highlighted. Sites that did not meet the filtering criteria are listed in **Additional File 2** and per-site counts of samples modified for gene/variant call-type pairs which passed the stoichiometric difference p-value filter are listed in **Additional File 3**

**Table 2.** Differential modification burden at the local level.

| Gene | Global | Type | AD | Position |
| --- | --- | --- | --- | --- |
| *ANXA11* | AD-enriched | A-G | 20 | chr10:80155380 |
| *CCDC66* | AD-enriched | A-G | 24 | chr3:56593020 |
| *DLG1* | AD-enriched | del | 21 | chr3:197043484 |
| *IFITM2* | CN-enriched | T-C | 17 | chr11:308290 |
| *MRPS21* | AD-enriched | T-C | 28 | chr1:150308112 |
| *PDZD2* | AD-enriched | del | 22 | chr5:32109039 |
| *SPECC1* | AD-enriched | T-G | 18 | chr17:20315480 |
| *TMEM130* | AD-enriched | del | 17 | chr7:98847504 |
| *ZNF84* | AD-enriched | T-C | 24 | chr12:133059027 |
|  |  |  | 27 | chr12:133062243 |
|  |  |  | 15 | chr12:133062765 |
|  |  | A-C | 29 | chr12:133057864 |

*Note:* Differential modification burden was assigned for each site passing filter criteria at the local level. Because all significant sites in this table were increased in AD, CN counts are zero for all sites and not shown. “Position” indicates the genomic position each modified site mapped to.

**Table 3.**
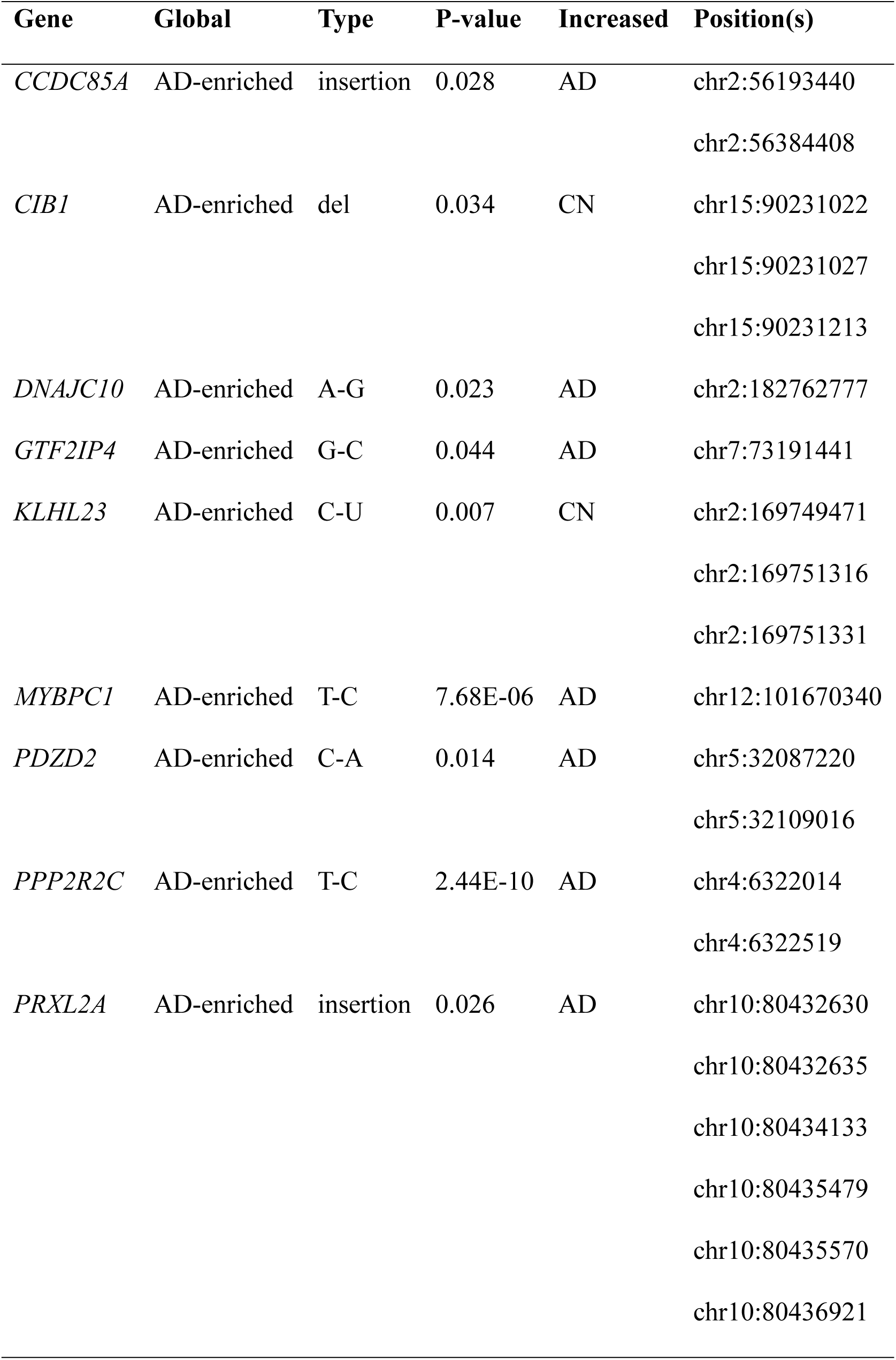
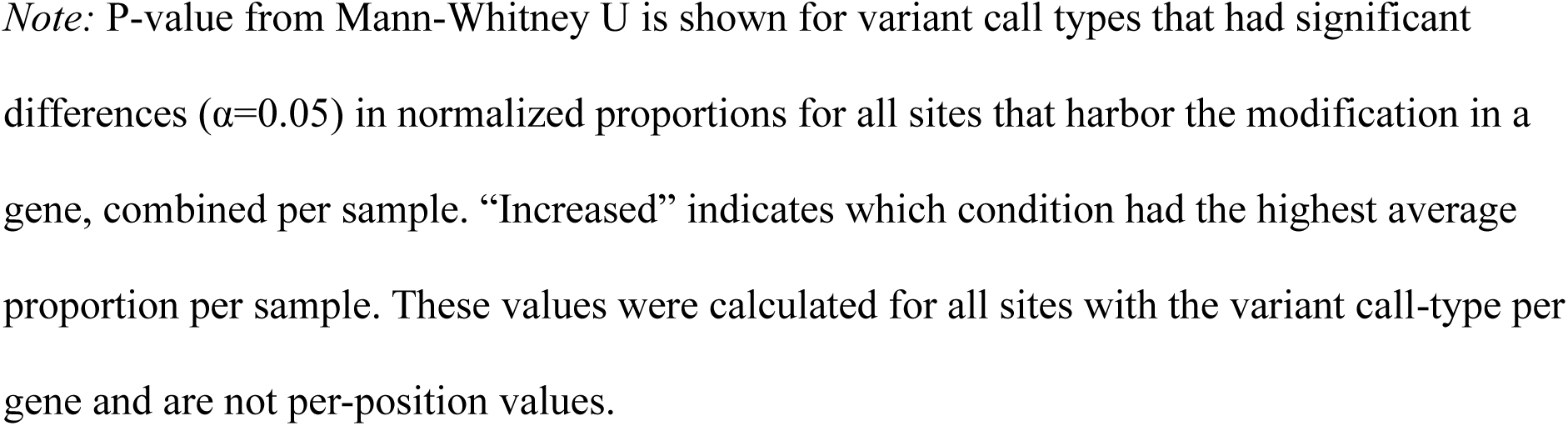
Differential modification stoichiometry by variant call-type at the local level.

| Gene | Global | Type | P-value | Increased | Position(s) |
| --- | --- | --- | --- | --- | --- |
| *CCDC85A* | AD-enriched | insertion | 0.028 | AD | chr2:56193440 |
|  |  |  |  |  | chr2:56384408 |
| *CIB1* | AD-enriched | del | 0.034 | CN | chr15:90231022 |
|  |  |  |  |  | chr15:90231027 |
|  |  |  |  |  | chr15:90231213 |
| *DNAJC10* | AD-enriched | A-G | 0.023 | AD | chr2:182762777 |
| *GTF2IP4* | AD-enriched | G-C | 0.044 | AD | chr7:73191441 |
| *KLHL23* | AD-enriched | C-U | 0.007 | CN | chr2:169749471 |
|  |  |  |  |  | chr2:169751316 |
|  |  |  |  |  | chr2:169751331 |
| *MYBPC1* | AD-enriched | T-C | 7.68E-06 | AD | chr12:101670340 |
| *PDZD2* | AD-enriched | C-A | 0.014 | AD | chr5:32087220 |
|  |  |  |  |  | chr5:32109016 |
| *PPP2R2C* | AD-enriched | T-C | 2.44E-10 | AD | chr4:6322014 |
|  |  |  |  |  | chr4:6322519 |
| *PRXL2A* | AD-enriched | insertion | 0.026 | AD | chr10:80432630 |
|  |  |  |  |  | chr10:80432635 |
|  |  |  |  |  | chr10:80434133 |
|  |  |  |  |  | chr10:80435479 |
|  |  |  |  |  | chr10:80435570 |
|  |  |  |  |  | chr10:80436921 |

*Note:* P-value from Mann-Whitney U is shown for variant call types that had significant differences ( $\alpha=0.05$ ) in normalized proportions for all sites that harbor the modification in a gene, combined per sample. “Increased” indicates which condition had the highest average proportion per sample. These values were calculated for all sites with the variant call-type per gene and are not per-position values.

Of the 17 genes found with the greatest local differences in either modification load or stoichiometry, 16 were AD-enriched, eight showed differential modification burden at one or more positions, and eight had differential stoichiometry of a variant call-type across all positions. Notably, one gene (*PDZD2*) appeared in both filtering lists, showing both differential modification burden at a single deletion site as well as differential stoichiometry for the C-A call-type. When per-sample events were compared, co-modification events were observed between genes and within genes. For the between-gene observations, each sample containing the *TMEM130* deletion also carried the *SPECC1* T-G variant. Most notably, however, was the within gene assessment of *ZNF84*: all 29 samples had the A-C variant call at a single, common position in addition to at least one T-C variant call at one or more position(s) across the four commonly modified positions highlighted for the gene.

## Discussion

### Modification signatures in RNA-seq datasets

When modification counts for all samples were summed by each variant call-type across all genes in the global set of 679 genes, we found that all variant call-types appeared in the AD group at a higher frequency than in the CN group, with most types also showing significant differences in modification stoichiometry of expressed transcripts. Data for both conditions were extracted and sequenced using the same stranded protocols and were likewise subject to identical data processing pipelines, reducing measurement and selection biases. Furthermore, by comparing counts and proportions between two conditions with similar age distributions, we aimed to reduce age and heterogeneity biases known to exist in epitranscriptomic analyses of age-related diseases, extracting only the patterns and sites that were notably different between AD and CN brains. Our analysis of variant calls from RNA-seq data used a combination of previously applied, suggested, and novel approaches designed to identify variants that are most likely to be products of post-transcriptional modifications and not errors arising from RNA processing or sequencing signal processing (32,51,52). From our results, we observed that the global distribution of variant call-types reflects previously published results (53,54).

One challenge to studying all modification types from transcriptomic data lies in the fact that, while current sequencing chemistries and technologies can sometimes differentiate single post-transcriptional modification types, none can confirm the original modification source for all variant call-types. This may be apparent in our findings that A-G (typically used to identify A-I edits) and C-T (typically used to identify C-U edits) variant call-types, along with their reverse complements (T-C and G-A), occur at a much higher frequency in both conditions than other single-nucleotide variant calls. Notably, these are the modifications for which existing mechanisms have been characterized, and they are generally highlighted in epitranscriptomic studies at the expense of other variant call-types: primarily because, as stated by Guo et al. (55), “HTS studies reported appreciable amounts on RNA variant call-types (yet)…whether non-canonical RNA editing events genuinely exist is an open question”, and that other post-transcriptional modifications are not easily identified from standard cDNA sequencing (54,56).

Considering these limitations, the study described here does not confirm that each variant call-type is a signature for a specific modification type but instead infers epitranscriptomic modification signals identifiable from analysis of stranded, short-read, RNA-seq data. A certain background rate is expected for false-positive variant calls in sequencing data, even after sequence cleanup and quality filtering; however, we expect those rates to be similar between the conditions and thus filtered during our comparative analysis. As seen in our global and local analysis results, we identified recurring modification sites of non-canonical variant call-types that differed significantly between genes and conditions. This, along with previous reports (57), suggests that these are not purely random variant calls. Together, the data support retention of all variant call-types in epitranscriptomic studies, investigating each equally, regardless of whether the chemical nature (e.g., editing, methylation, pseudouridylation) of the mismatched nucleotide has been experimentally determined.

### A novel look into epitranscriptomic regulation in aging and AD

To our knowledge, this study provides the first in-depth analysis of epitranscriptomic changes in AD and aging from RNA-seq data. Our results highlight gene subsets that were hyper- or hypo-modified in AD, global and local distributions of recurring and sporadic modification sites, pathways susceptible to epitranscriptomic regulation, and differences between conditions based on both counts (modification burden) and proportions (stoichiometry of modification occupancy).

#### Sporadic and recurring modification sites

The literature shows that epitranscriptomic modification patterns are context-dependent and that they can be widely disrupted in disease (22,23,58). The heterogeneous nature of the epitranscriptome can complicate analyses, so we decided to filter through the noise, assessing the distribution of recurring (modified in >1 sample) and sporadic (modified in exactly one sample) modification sites. We observed a global increase in sporadically modified sites in the brains of AD samples. When assessed individually, the single sites are difficult to interpret, yet they collectively reveal a pattern of hyper-modification in the AD condition. Additionally, many sites that were identified as recurring were shared between conditions, yet our local level analysis identified sites specifically recurring only in AD, with 50% to all of AD samples harboring the modified site while none of the CN samples did. We hypothesize that the recurring sites shared between conditions are likely related to normal aging, whereas recurring sites within a single condition are indicative of condition-specific changes. Sporadic modifications, especially when increased in AD, suggest altered modification enzymes or pathways in AD, or other ‘omics level changes which may lead to aberrant signaling for modification of transcriptomic sites in a seemingly random approach (23,59,60).

#### AD-enriched pathways

To investigate pathways most likely under epitranscriptomic regulation in the aging brain, we conducted a pathway analysis on the genes found to be hypermodified in AD. Among the significantly-enriched pathways identified were “protein processing in the endoplasmic reticulum,” “renal cell carcinoma,” “chronic myeloid leukemia,” “Kaposi sarcoma-associated herpesvirus infection,” and “neurotrophin signaling pathway.” The “protein processing in the endoplasmic reticulum” pathway is often associated with AD due to the nature of the disease as a proteinopathy, characterized by the accumulation of misfolded Tau and Aβ (61,62). The connection between AD and “renal cell carcinoma” or “chronic myeloid leukemia” is less clear but may be due to inflammation as a shared pathologic feature of the diseases (63) or may point to similar accumulation of DNA damage (64). Intriguingly, the Kaposi’s sarcoma-related herpesvirus (KSHV) infection pathway, which has been indicated as one of the most influential infection-related pathways to act on other AD-related pathways (65), was in the top five enriched in our study and showed the highest number of gene overlaps with the other highly enriched pathways.

#### Showcase of most significant differences at the local level by position and variant call-type

We wanted to home in on local differences between the epitranscriptomes of AD and cognitively normal brains, so after filtering on aggregated proportions by variant call-type, we assessed variant call-type proportions and per-position counts from 24 genes, showcasing a final set of 17 genes with either differential modification burden by position or significantly different variant call-type stoichiometry. Several of these genes, such as *ANXA11* (66–68), *IFITM2* (69,70) and *PDZD2* (71–75), have been implicated in neurodegeneration, neuroinflammation, and/or as viral targets during infection. Our analysis provides a new potential mechanism through which these gene transcripts may be regulated to impact disease processes.

Interestingly, our extended protein-protein interaction search found that protein products of four genes out of the showcased 17, *SPECC1*, *ZNF84*, *MRPS21* and *PP2R2C*, were identified as potential interaction partners with other proteins involved in the KSHV infection pathway, either directly or indirectly via *GNB1* and /or *GNB4.* Neither of the *GNB* genes were identified as significant during our local level analysis but were highlighted by Enrichr-KG, from our list of AD-enriched genes, as contributing to the KSHV infection pathway. An additional search of the literature identified *CIB1* as important in both KSHV and AD (76–79).

Most of the research on KSHV (HHV-8) focuses on its role in cancer development and other diseases, but it has been established that cancer and AD share an inverse comorbidity. This suggests that a single event acting on the human system can either deregulate processes in a way that leads to cancer or AD, but rarely both (80–84). Although viral status is not known for the samples in this study, it is believed that HHV-8 has neuroinvasive capabilities where it can reside silently in neural tissue without presenting the typical symptoms associated with the virus, and it may infect a much larger portion of the population than is often reported (85). Recent work has shown that KSHV (and other herpesviruses) extensively rewires the host epitranscriptome upon entering the lytic cycle, altering the pathways that control lytic replication by removal and addition of modifications, strategically silencing host defenses or hijacking useful host molecules (86). Furthermore, viral infection has been repeatedly linked to neuroinflammation and AD, and the inclusion of this pathway in our most significant results re-emphasizes the potential connection between viral infection and epitranscriptomic alterations in AD (87–90).

The observations made in this study reflect the epitranscriptomic features that would be expected of herpesvirus infection in the brain. These include a drastic change in both sporadic and recurring modification sites where many are found in critical genes involved in host immune response and defense. Future investigation of these connections and how they may lead to a cascade of deregulation onto other pathways could provide exciting new links to the multi-omics network (71,91) upon which disease outcomes depend.

### Conclusion

The results from the current study provide a targeted baseline of transcripts and nucleotide positions as high priority targets for additional investigation. These data support the inclusion of all variant call-types, and both global and local stratification analyses, in future large-scale epitranscriptomic assessment algorithms. Future studies may reveal the biologic and pathogenic significance of these modifications and point to novel biomarkers or precision medicine approaches for AD therapeutics.

## Declarations

### Ethics approval and consent to participate

“The parent studies and sub-studies were all approved by an Institutional Review Board of Rush University Medical Center and all participants signed an informed consent, Anatomical Gift Act, and a repository consent to share data and biospecimens.” (92)

### Consent for publication

Not applicable.

### Availability of data and materials

The results published here are in whole or in part based on data obtained from the AD Knowledge Portal (93). Study data were provided by the Rush Alzheimer’s Disease Center (RADC), Rush University Medical Center, Chicago. Data collection was supported through funding by NIA grants P30AG10161, R01AG15819, R01AG36836, U01AG32984, U01AG61356, P30AG072975, the Illinois Department of Public Health, and the Translational Genomics Research Institute (genomic).

The datasets supporting the conclusions of this article are protected and not publicly available due to participant privacy and the terms of the Data Use Agreement. However, they are available to qualified researchers upon request to AD Knowledge Portal (93).

The custom code and scripts created for use in this study are available on GitHub (94).

### Author’s contributions

Katie Jensen is credited with Methodology, Formal Analysis, Investigation, Writing Original Draft and Editing, and Data Visualization. Jennifer Weller is credited with Review and Editing of the Manuscript and Project Supervision. Kristen Funk is credited with Project Conceptualization, Providing Resources, Writing Original Draft and Editing, and Funding Acquisition. All authors read and approved the final manuscript.

## Acknowledgements

The data available in the AD Knowledge Portal would not be possible without the participation of research volunteers and the contribution of data by collaborating researchers. We thank Dr. Jessica Schlueter and Dr. Anthony Fodor (UNC Charlotte, Bioinformatics and Genomics) for guidance in data analysis and careful reading. We appreciate the contribution Cynthia Gibas (UNC Charlotte, Bioinformatics and Genomics) to the graphical abstract concept. We also thank Jack Young (UNC Charlotte, Bioinformatics and Genomics) for his valuable suggestions concerning parallel processing and algorithm optimization.

## Additional Files

### 1. Additional File 1

*AdditionalFile1.csv*

Table of counts for distribution of unique and occurring modifications by variant call-type, gene category within which a modification is identified, and the odds-ratio and p-value outcomes from a Fisher Exact test of the counts. This file was used to create the Figure 4 forest plot.

### 2. Additional File 2

*AdditionalFile2.csv*

Table of modified sites from local-level top 24 gene set not included in local-level 17 gene set. List of all modified sites from top 24 gene set which did not pass the final modification burden nor occupancy filters. Sample counts are per site, stoichiometric values are from Mann-Whitney U of average normalized proportions per sample for all modifications of variant call-type.

### 3. Additional File 3

*AdditionalFile3.csv*

Table of sample counts from modified sites of local-level top 17 genes.

List of sample counts at all modified sites in top 17 genes for (gene, variant call-type) pairs with significant differences in stoichiometry at α=0.05. The condition columns indicate the number of samples in the condition that harbor the modified site.

## List of Abbreviations

AD: Alzheimer’s Disease
BAM: binary alignment map
CN: Cognitively Normal
EOFAD: Early Onset Familial AD
FASTA: nucleotide sequence files
FASTQ: sequencing data files
GTF: general transfer format files
KSHV/HHV-8: Kaposi’s sarcoma-associated herpesvirus
LOAD: Late Onset AD
PONs: panel of normals
RNA-seq: RNA sequencing
VCF: variant call format file
WGS: whole-genome sequencing

